# Active modules for multilayer weighted gene co-expression networks: a continuous optimization approach

**DOI:** 10.1101/056952

**Authors:** Dong Li, Shan He

**Affiliations:** School of Computer Science, The University of Birmingham, UK Zhisong Pan, Guyu Hu; PLA University of Science and Technology, China

## Abstract

**Motivation:** Searching for active connected subgraphs in biological networks has shown important to identifying functional modules. Most existing active modules identification methods need both network structural information and gene activity measures, typically requiring prior knowledge database and high-throughput data. As a pure data-driven gene network, weighted gene co-expression network (WGCN) could be constructed only from expression profile. Searching for modules on WGCN thus has potential values. While traditional clustering based modules detection on WGCN method covers all genes, unavoidable introducing many uninformative ones when annotating modules. We need to find more accurate part of them.

**Results:** We propose a fine-grained method to identify <u>a</u>ctive <u>mo</u>dules on the m<u>u</u>lti-layer weighted (co-expression gene) <u>n</u>e<u>t</u>work, based on <u>a</u> cont<u>in</u>uous optimization approach (AMOUNTAIN). The multilayer network are also considered under the unified framework, as a natural extension to single layer network case. The effectiveness is validated on both synthetic data and real-world data. And the software is provided as a user-friendly R package.

**Availability:** Available at https://github.com/fairmiracle/AMOUNTAIN

**Supplementary information:** Supplementary data are available at *Bioinformatics* online.

## 1 Introduction

As a well-known fact, a group of genes may get involved into a biological process other than act alone [3], thus identifying a group of genes and associating it with certain biological functions is of important. In this paper, we define a **functional module** in a biological network as a subnetwork which may involve a common function in biological processes. Another important but different concept is **topological module**, may also be referred as community, within which the interactions are much more intensive compared with those outside [15]. Topological modules have been studied intensively and the modular structure is easy to be detected in general sense, but functional modules are of real interest [3].

Although a function module may overlap with a topological module, only using the network structural information is not enough to find the function modules. The topology of a biological network does not always precisely reflects the function or even disease-determined regions [3], which are the real concerns in biology. To bridge the gap between the topological module and functional module, searching for **active modules**, i.e. connected regions of the molecular interaction network showing striking changes in molecular activity or phenotypic signatures that are associated with a given cellular response [28], has become a central challenge in system biology. Active modules were shown to be able to reveal regulatory mechanisms [19], which closely to function modules, i.e., these modules might connect multiple function modules. The activities of network nodes are usually measured by high-throughput omics data. In recent years, many active module identification algorithms have been developed to solve this problem, and most of them are applied on a *skeleton networks* plus *muscle* paradigm. The skeleton networks like protein-protein interaction networks or metabolic networks are constructed from prior knowledge database [19, 27, 10]. However, compared with increasing vast amount of high-throughput omics data, the speed of constructing reliable and complete skeleton networks, which heavily rely on experiment and human validation, is quite slow. For some non-model species, or even some new model species such as Daphnia, the PPI network even does not exist. The lack of reliable skeleton networks posts a challenge to the detection of active module to reveal certain key mechanisms in the biological systems.

In contrast, gene co-expression network is a pure data-driven gene network, which could be constructed only from expression profile. In such networks a nodes is a single gene and an edge is the correlation relationship between a pair of genes. And a weighted gene co-expression network is a fully connected graph. Modules in such networks are also considered to participate into some biological process [44], and those modules with significant biological meaning are essentially functional modules.

As the first but crucial step for modules functional annotation analysis, module identification on gene networks is an important but less studied topic. Traditional module detection on gene co-expression networks was based on gene clustering, i.e. putting similar genes based on their correlations or edge weights into clusters as modules [44]. The coarse-grained clustering technique basically covers all genes in the network. As a results, the identified functional module including those genes show very little activities, which might not be very informative to reveal the biological mechanisms. We hypothesize that by identifying active modules which consider gene activities in the coexpression network, more precise biological mechanisms would be obtained. How to rigorously define **active modules** in such weighted network is still an open problem, but the module itself should be be more compact and informative compared with random subnetwork or clusters from two perspectives: 1) From the topological view, active modules are supposed enjoy high module scores measured by nodes and edges. 2) From the functional view, active modules are supposed to be more significantly associated with some biological process. The active modules identification on gene co-expression network is also a new problem, especially for the multilayer gene networks. A better understanding of modules in such network is expected to establish a pipeline from pervasive gene expression data to reasonable biological interpretation on a systemic level.

As a generalization of the single network, there are various reasons to model the interactions in living organisms as multilayer networks. Different layers may represent different time points, multiple conditions or various species. Modules across multiple layers in this report may reveal some properties of weighted gene co-expression networks such as time-invariant component genes, general responsive functional modules, and species conservation biological process. Similar but more general topics are also called multiplex networks [29, 22]. An existing work [24] mined recurrent heavy subgraphs in multi-slice networks, where each network shares the same set of genes without interactions between them. Conversed modules identification examples include [48, 13] which will be mentioned later. Inspired by [24], we develop a unified optimization framework to identify active modules on the multi-layer weighted co-expression gene network (AMOUNTAIN), and the layers could cover all three cases mentioned above.

## 2 Methods

In the early settings, active module identification was proposed to find significantly changed subnetwork in modular interaction networks [19]. Most following works developed methods based on the “skeleton+muscle” paradigm, where the “skeleton” is basic molecular interaction network constructed from prior knowledge database, and the “muscle” comes from the widely available high-throughput data which measure the genes activities. In general the skeleton biological network is represented as an undirected graph *G* = (*V*, *E*), where nodes in *V* represents components like genes (or gene products), proteins or metabolites, and edges in *E* represents the interaction between two nodes. Each node *i* is assigned a single score to denote the activity of corresponding component in certain condition, such as fold-change or p-value of gene expression level. The simplified problem of finding highest score module in unweighted network, which consider the subnetwork score is the sum of each node’s score, is formally defined as following:

#### problem 1

*Given a graph G* = (*V,E*) *with vertices weights* **z** ϵ *R_n_* for each *v* ϵ *V find a connected subnetwork S* = (*V_S_, E_S_) of G with maximal weight* 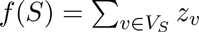

This combinatorial optimization problem is also called Maximum-Weight Connected Subgraph Problem (MWCSP), which is equivalent to finding a maximum weight clique in a weighted graph, being referred as a famous NP-complete problem [20]. The proof is provided as supplementary materials of Ideker et al. [19]. As effective tools to solve combinatorial problems, metaheuristic algorithms have been widely applied to search satisfied solutions. The original paper [19] proposed to use simulated annealing, a generic probabilistic metaheuristic to solve this problem. Other methods include extended simulated annealing [17], greedy algorithm [40, 41], graph-based heuristic algorithm [34], genetic algorithm [27] and some exact approaches based on integer linear programming [33, 10, 46, 2].

### 2.1 Single-layer network

Compared with increasing vast amount of high-throughput data, the speed of constructing or confirming precise molecular interactions, which heavily rely on experiment and human validation, is quite slow. With only gene expression data, we could build gene co-expression network (GCN). We generalize the idea of active modules in problem (1) to GCN case, and consider the node score as gene importance criterion in gene co-expression network, which can be evaluated by the expression level changes (such as fold change or other more comprehensive statistics) under certain conditions. Meanwhile the network structure is also determined by gene expression data and expressed as a weighted network. In practice, we get a bit more different problem to find a subgraph of size k (otherwise it corresponds to a trivial case containing all nodes) which aims to has both maximal node weights and closely connected to has large edge weights, formally defined as:

#### problem 2

*Given a complete graph G* = (*V,E), with vertex weight z_v_* ϵ *R for each v* ϵ *V and edge weights W* = [*w_ij_*] *for each edge* (*i*,*j*), *find a subgraph T of size k with large vertices weight* 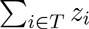 *and also edges weights*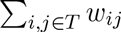.

Problem (2) is actually a simplified problem of (*K*_1_, *K*_2_)-Recurrent Heavy Subgraph (RHS) problem [24] but with additional node scores. (*K*_1_,*K*_2_)- RHS considers multiple co-expression networks, which is also discussed in details at next section. The module can be represented by membership vector x ϵ {0,1}^n^, where *x*_i_ = 1 means *i*-gene belongs to the module. Thus the optimization is naturally expressed as:

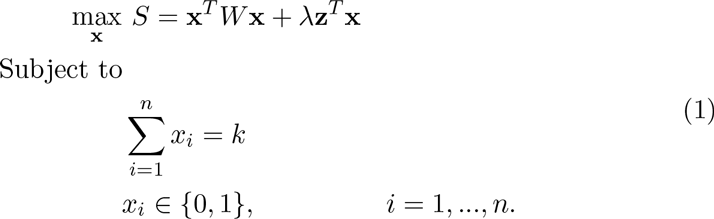

The NP-hardness can be proved by reducing the well-known NP-complete problem *k*-clique to this problem, for the details of the proof refer to the supplementary materials. Although integer programming methods [33, 10, 46] can be applied, it may cause high computational complexity and be lack of theoretical guarantee w.r.t running time and accuracy. Alternatively, if we relax the integer constraints of x to continuous constraints [42, 24] and control the module size by introducing a vector norms of **x**, it becomes a nonnegative and equality constrained quadratic programming (QP) problem (2), which can be solved by various existing continuous optimization techniques in polynomial time.

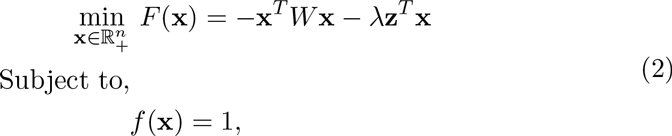

where *f* (x) is the vector norm. The *l_p_*-norm (*p* > 0) of x is defined as
(∑ _i_ |*x_i_*|^*p*^)^1*/p*^.

The choice of vector norm has an impact on how to solve the problem (2). For example, the 4)-norm can produce a sparse solution which is consistent with the fact that only a few members belong to the module. The *l*_0_-norm is widely used as an alternative to *l*_0_ since the optimal solution of the latter corresponds to a combinatorial problem [11]. The *l*_2_-norm is also widely used for closed-form solution but cannot lead to a sparse solution. A linear combination of *l*_1_ and *l*_2_, i.e. (1 — α)‖x‖1 + α‖x‖^2^ is also called elastic net penalty [49], when the objective is least square and α = 0 corresponds to lasso [38] and α = 1 corresponds to ridge regression. Elastic net is considered to enjoy the characteristics of both lasso and ridge regression. Besides, the *l*_∞_-norm is *max*{*x*_1_, *x*_2_,…, *x_n_*} which makes the values in vector smooth and all the entries are roughly identical. All the mentioned vector norms and the existing corresponding optimization techniques may be applied in the problem (2) as long as the constraints of vector norms can reveal some essence of module membership in weighted co-expression network. While the fundamental one is *l*_1_-norm since the module size needs to be constrained, otherwise, the problem becomes trivial with all nodes included in the target module.

The *l*_1_-norm constraint of optimization problem (2) can be converted to 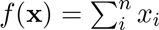 since *x_i_*≥ 0. This problem has been intensively studied in mathematical programming [7, 4] and could be solved by a lot of existing method. But only using *l*_1_-norm constraint tends to produce very sparse solution for problem (2), even a vector x which contains only one non-zero element may be the optimal solution. [24] used the mixed norm *l*_0,∞_,(x) = =‖x‖_0_ + (1 — α)‖x‖∞ (0 < α < 1) to encode the characteristics of gene membership, where the optimal vector should contain equal non-zero values and the rest zero values. In practise *l*_0,∞_, was approximated by *l_p_,_2_*(*x*) = α‖*x*‖_*p*_ + (1 — α)‖x‖_2_ (0 < *p* < 1), and they solved the non-convex problem (1) with only edge weights by MultiStage Convex Relaxation (MSCR) [45].

It is a natural idea to use the elastic net penalty [49] to control the sparsity and achieve desirable membership, i.e. *f (x)* = (1 — α)|x|_1_ + α‖x‖^2^. And a general strategy follows the gradient projection method [25] and generate a sequence to approximate the accurate solution like

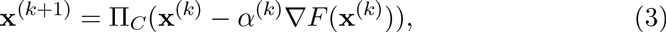

where ∏_C_ is the Euclidean projection of a vector on convex set *C*, defined as (4):

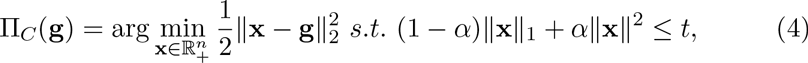

where **g** is a constant vector and *t* is the radius which has little impact on the solution in practice. The step size α^(*k*)^ in *k*-step should satisfy the following condition (5) in order to make the objective function:

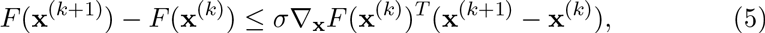

where σ is a small positive constant. Searching for optimal α^(*k*)^ is time consuming. Here we adopt the same as in [25] that scale α^(*k*)^ by a fixed factor β until α^(*k*)^ satisfies (5). Thus the algorithm is guaranteed to converge.

Solving subproblem (4) involves a root finding procedure [16] which can be done in linear time, and the total iterative procedure can be improved up by Nesterov’s method [30], which replace the current step **x**^(*k*)^ in (3) with a linear combination of previous two steps, **s**^(*k*)^ = **x**^(*k*)^ + *t_k_*(**x**^(*k*)^ — **x**^(*k*—1)^) where *t_k_* is another parameter to make it convergence. Nesterov’s method has been shown to have optimal convergence rate for first-order method. Refer to the supplementary material the details about how to solve the convex optimization problem (2) with Elastic net penalty.

**Algorithm 1.**
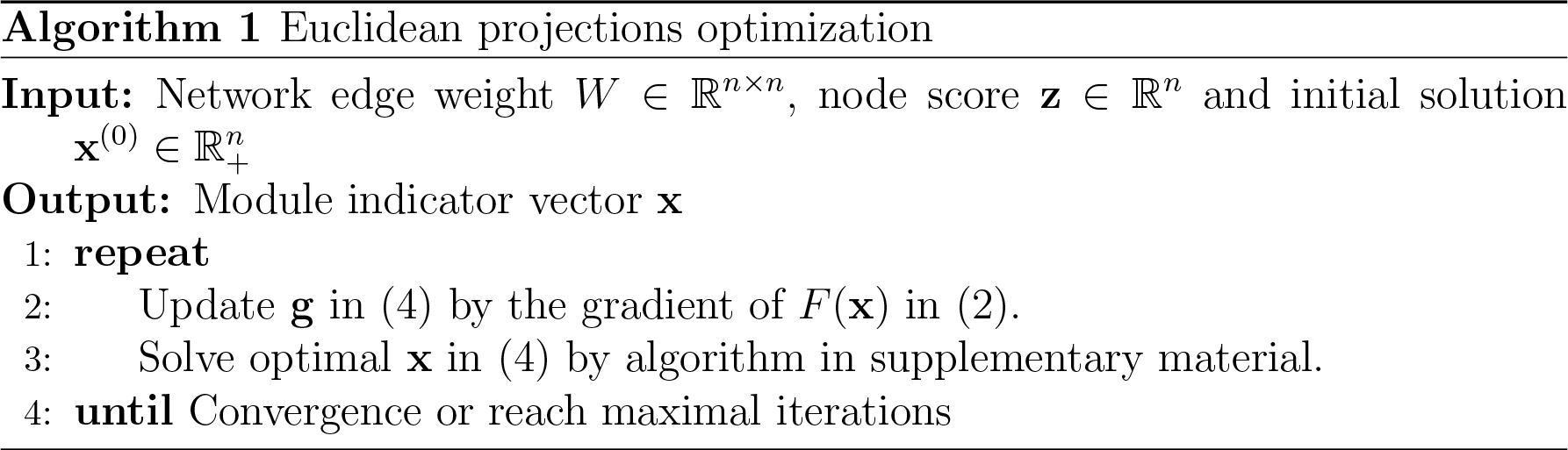
Euclidean projections optimization

Generally we may want to identify multiple modules from one network. Similar to [47, 26], we can find *N* modules by running algorithm 1 *N* times, with each time simply extracting the resulted module from background network. The resulting sequences of modules may indicates the importance in terms of node activities and correlation similarities, in a descending order. The general procedure for identifying *N* modules from given gene expression profile can be summarized as algorithm 2.

### 2.2 Two-layer network

The biological system has been modeled as a multi-layer network before [5], while more detailed analysis on multi-layer networks raises up in recent years [21], after the intensive research on single layer networks. The multi-layer network provides a general framework to model temporal and spatial change of interactions for cellular networks, and contributing three aspects for current computational biology research: 1) Modeling dynamic properties for biological process as multiple snapshots, 2) Modeling different responses to multiple conditions of the same species and 3) The identification of conserved genes across species as well as specific genes.

**Algorithm 2.**
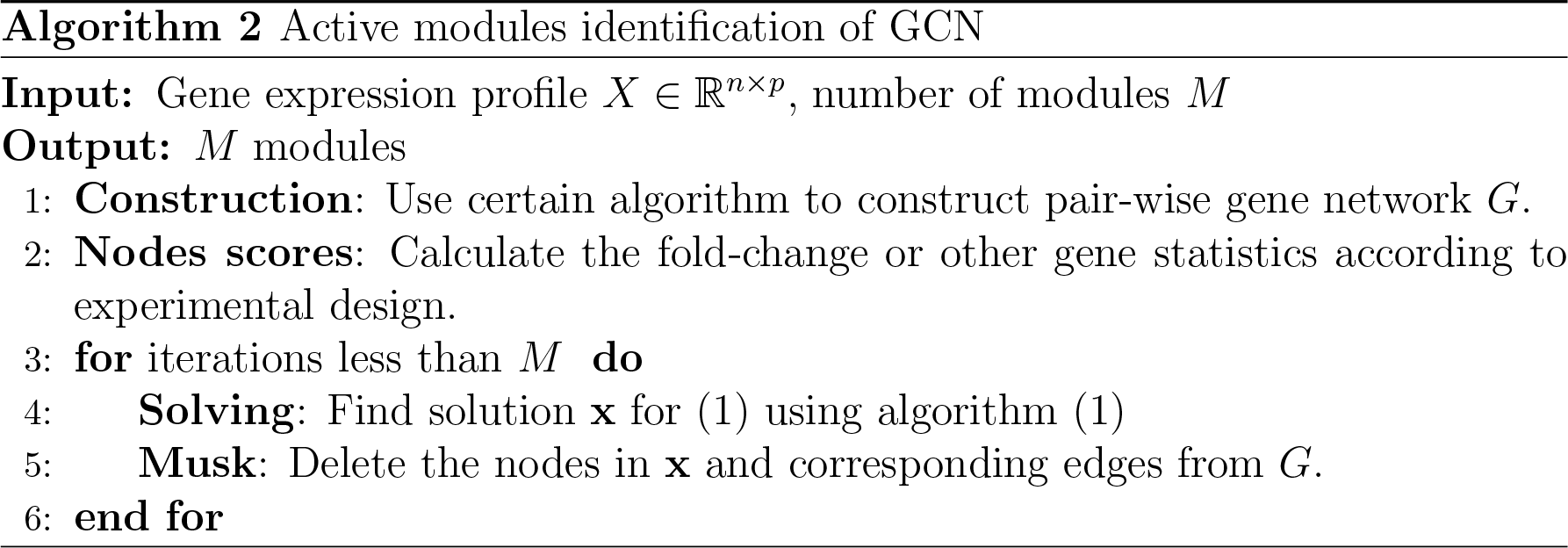
Euclidean projections optimization

Although the behaviors of living organisms were considered to be dynamic, traditional network-based methods primarily focused on static network, which is a snapshot of the real case. Multi-layer networks provide a powerful tool for modeling this time series networks, with each layer standing for a time point. The differential analysis on this time series networks may reveal several important concepts related to time changing on cells belongs to certain species or tissues, such as the key components that were not be affected or some vanishing structures.

The integration of genomic techniques into environmental toxicology has shown potential application values to develop exposure biomarkers and investigate the mode of toxicity [31]. Representing the interaction of a set of genes’ response under different conditions as multiple layer network may shed light no this evolutionary conservation of living organism.

Besides the dynamic change of networks in the same set of components, a multi-layer network can also capture the core set across species, such as conserved active modules [37, 9]. By constructing a multi-layer network with each layer representing one species, we may find similar patterns in a species that has relatively more prior information, thus to gain biological knowledge of interested species. Finding conserved modules may also improve our understanding about the evolutionary biological procedure by highlighting the similarities and differences in key patterns between species [48].

Although [24] was proposed for multiple gene co-expression networks, which is similar to multi-layer networks. They can be distinguished from two aspects: 1)Multiple networks share exactly the same set of nodes while multi-layer networks do not necessarily to, which makes later can be applied in multiple species orthology. 2)Multi-layer networks consider inter-layer interactions while multiple networks do not to.

Another related work is `xHeinz` [13], which mines cross-species network modules using an integer linear programming approach. It extends the single network module identification algorithm `heinz` [10] to two species case and takes the same optimization technique for the problem. As discussed before, this kind of algorithm requires the skeleton interaction network plus high-throughput data, which is the main difference between our work. The network node scoring function in `xHeinz` is inherited from that in `heinz`, which requires the parameters estimation in a beta-uniform mixture (BUM) model. While our algorithm simply uses the fold-change information since node score is only part of the whole objective. In theory we can also use the adjusted log-likelihood ratio score in `heinz`. The optimization methodology we adopt is also straightforward, while `xHeinz` relies on external integer programming solver CPLEX.

Being similar to single layer network situation, an active module in a two layers-network can be represented as a connection of two modules in two different networks *G*_1_ = (*V*_1_,*E*_1_) and *G*_2_ = (*V*_2_,*E*_2_). The inter-layer interactions were measured by *A* = *[a]_i,j_* ϵ ℝ^n_1_×n_2_^ where *n_1_* and *n_2_* are the numbers of nodes in *G*_1_ and *G*_2_. The basic two layer network module identification problem is formally defined as

#### problem 3

*Given two complete graphs G*_1_ = (*V*_1_,*E*_1_) *and G*_2_ = (*V*_2_,*E*_2_), *with vertices weights z_1 v_* ϵ *R for each v* ϵ *V*_1_ *and z*_2*v*_ ϵ *R for each v* ϵ *V*_2_. *And edges weights W*_1_ *for edges in G*_1_ *and W*_2_ *for edges in G*_2_. *The interlayer interactions were measured by A* = [*a*]_*ij*_ ϵ ℝ^*n*_1_×*n*_2_^. *The goal is to find two subgraphs T*_1_ ϵ *G*_1_ *and T*2 e *G*_2_ *which both have large vertices weights and edges weights as well as intensive interaction with each other.*

We use two two variables **x** and **y** to represent the memberships of active modules in two different networks, *x_i_* = 1 means the *i*-th node in the first network is in the module. Thus the optimization problem can be expressed as an extension to (2),

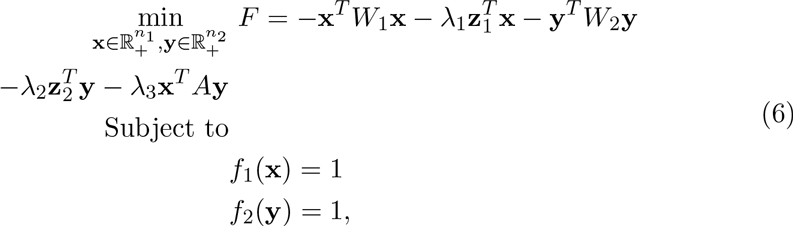

where *f*_1_(*x*) and *f*_2_(*y*) are the vector norms on two vectors respectively. For simplicity we use the same Elastic net penalty 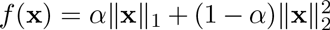 or mix norm penalty *f*(*x*) = α‖x‖_*p*_ + (1 — α)‖x‖_2_ for both **x** and **y**. The general idea for solving (6) is alternating optimization, i.e. a iteratively optimize one variable while fixing another each time [25]. When optimizing one variable, it has the same form as in (2). Each iteration in the procedure can be simply expressed as:

- Find x^(k^+^1)^ such that *F*(*x*^(*k*+1)^, *y*^(*k*)^) ≤ *F*(*x*^(*k*)^, *y*^(*k*)^) and,
- Find *y*(^*k*+1^) such that *F*(*x*^(*k*+1)^, y^(k^+^1)^) ≤ *F*(*x*^(*k*+1)^, *y*^(*k*)^)

The complete algorithm to find multiple modules in the two-layer network shares the same structure of algorithm 2. There is another parameter λ_3_ in (6) controlling how much degree the inter-layer links affect the resulting modules. Take multi-species for example, large λ_3_ can leads to conserved modules across different species which may reveal some gene conservation in response to certain changes. Conversely, small λ_3_, e.g λ_3_ = 0 makes the interlayer information playing no role, thus leading to two independent module identification processes. If we have multiple layers more than two, the rational keeps the same. As long as each layer has different set of nodes, alternating optimization can be used as the same way as in two-layer networks. Otherwise a more compact tensor computational paradigm [24] can be more efficient without inter-layer links consideration.

## 3 Results

### 3.1 Synthetic data

Several related works have used artificially generated data [34, 40, 23, 33] in order to test their algorithms in single network module identification. Being different from previous networks, the simulated networks here should have clear topological structure as well as node scores. We follow [24] to construct gene co-expression networks for simulation study. Let *n* be the number of genes, and edge weights as well as node score follow the uniform distribution in range [θ,1]. A module contains *k* genes inside which the edge weights as well as node score follow the uniform distribution in range [9, 1], where θ={0.5, 0.6, 0.7, 0.8, 0.9}. Figure 1 shows the weighted co-expression network when *n* = 100, *k* = 20 and red nodes indicate module members and wider edges mean larger similarities. Visualization is based on `qgraph` [14].

With ground-truth in hand, we can define the following performance measurement for the problem (2). From the topological view, the *accuracy* for each single module identification is considered as a binary classification problem. The performance is defined by the 2 × 2 confusion matrix, including the number of genes correctly detected (True positive, *tp*), the number of genes in identified module but not in the real module (False positive, *fp*), the number of genes in the real module but not in the identified module (False negative, *fn*), and the number of genes neither in identified module or in the real module (True negative, *tn*).

**Figure 1:**
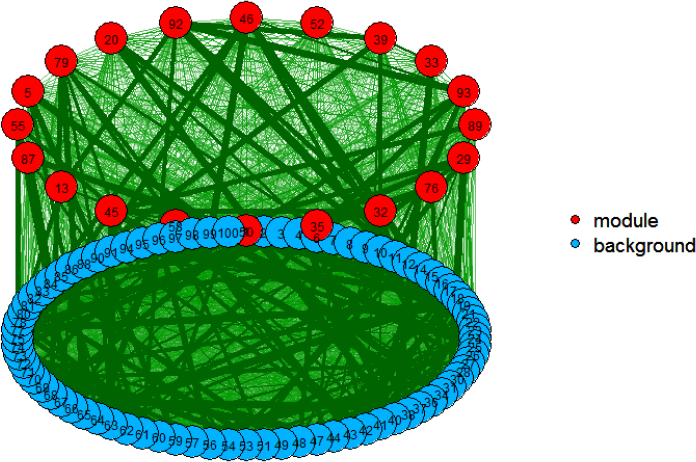
Simulated weighted gene co-expression network.

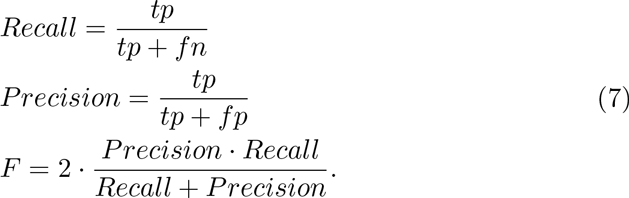

We conduct the simulation study on a relatively large network when *n* = 10000, and we consider a sparse case where one module contains *k* = 100 genes and node score follow the uniform distribution in range [θ,1], where θ= 0.5. The algorithm which uses mixed norm in (2) is outperformed by our algorithm 1 using elastic net penalty, in terms of both the running time and predictive accuracy. By choosing proper parameters from grid search combining α={0.1 ∼ 0.9} in elastic net penalty and λ=2^{-5^∼^5}^ in (2), we can almost exactly find the target model nodes. The optimal α={0.3 ∼ 0.4} and λ = 2^{-5^∼^—1}^ for this network which makes *F* = 1. Figure 2 shows how these parameters affect F-score (7) in this case. A similar result is also found in two-layer network simulation.

**Figure 2:**
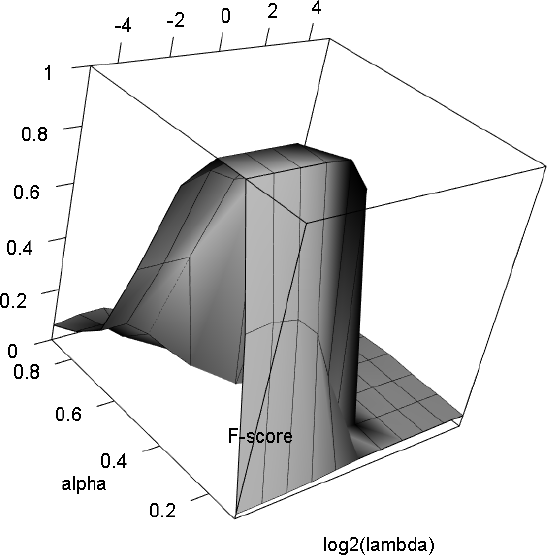
Parameters selection for algorithm 1 on large network.

### 3.2 Real-world data for single layer network

We mainly use gene expression datasets from Gene Expression Omnibus (GEO) [12] as real-world data examples. GSE3635 and GSEGSE5283 are two expression profiling by array from Saccharomyces cerevisiae. The original goal of the dataset is study the the regulation of transcription factor YOX1 and YHP1 during the cell cycle of Saccharomyces cerevisiae [32], with deleted YOX1 and YHP1 deleted. The wild-type and mutant cells were collected at 0, 10, 20, 30, 40, 50, 60, 70, 80, 90, 100, 110, and 120 minutes after synchronization with alpha factor. We take the formal as control group and second as experimental group since they share the same set of genes. And we assume the biological processes with YOX1 and YHP1 deleted was measured by gene expression values. The goal is to find several set of genes, referred as modules identification using algorithm 1, which are closely related to these biological processes. The gene expression values has been normalized as log ratio using Rosetta Resolver, we only need to deal with missing or invalid values. The strategy is simple, discarding probes with more than 20% missing values or NAs and repalcing missing or NAs positions in a valid probe with mean value of the rest samples of that probe. We donot pick up signigicantly expressed genes using linear model like many other methods, since the algorithm requres as complete information about gene correlation and gene activities only contribute part of the whole objective.

First, algorithm 1 requires the a weighted gene coexpression network as input. We employ the same differential analysis method of gene pairs as [18]. The difference of coexpression, also the edge score of gene i and j between two conditions *a* (control) and *b* (experimental) was quantified by,

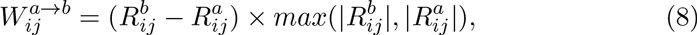

where 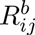 is the correlation between gene *i* and *j* under condition *b*. The node score which reflects the gene expression activity degree is measured by the ratio of expression level under two conditions,

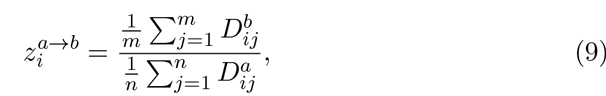

where 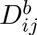 is the gene expression value of gene *i* in *j*-th sample under condition *b*, and *m*, *n* are the number of sample in condition b and a respectively. Here we let *N* = 10 in algorithm. The parameter λ represents the trade-off between edges weight and nodes weight which seems to play a slight role in performance. Here we simply fix λ = 1 and use a binary search method to select α for elastic net penalty which controls the sparsity of the module. See supplementary text section 4 for usage of parameter selection. Here we desire each module size with around 100 200 genes. These modules are provided as gene list (S1.xlsx).

We performed functional enrichment analysis of these modules. The basic of functional enrichment of a module is to assign the biological process annotations in Gene Ontology [1] to the genes (proteins) in that module. The probability that a module of size n have the same function as an existing functional module can be calculated by a hypergeometric distribution with Gene Ontology Term Finder. The P-value is calculated by the following formula,

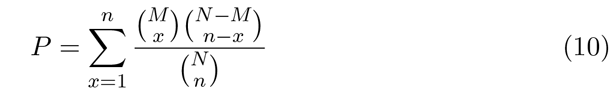

where *N* is the total number of genes (proteins) within the genome and M is the total number of genes (proteins) within a category. A low p-value indicates genes have high overlap with enriched functional categories thus are biologically significant. Results show that module identification using algorithm 1 can find consistent biological processes only from expression data which help to understand the underlying mechanisms related to biological conditions. Since YOX1 and YHP1 are important transcription factors in the regulation of the cell cycle, the identified modules are enriched by corresponding biological processes such as single-organism cellular process (G0:0044763), cellular macromolecule metabolic process (G0:0006139) and nucleobase-containing compound metabolic process (G0:0044260). Furthermore, the enriched GO terms are less significant as they are later identified, which corresponds with the algorithmic settings.

### 3.3 Real-world data for two-layer network

Inspired by xHeinz [13], we chose two expression data for two species mus musculus and homo sapiens: GSE43955 and GSE35103 for multilayer case. The original studies [43] and [39] reported the expression profiles identification controlled by the differentiation of Th17 cell. Here we expect the proposed algorithm could find consistent results from a two-layer gene co-expression network. Each layer of the network is constructed from the gene expression of a species, and the edge weights and node weights are defined by (8) and (9) as well. In each layer the two conditions are with or without Th17. Specifically, we use the expression value of two conditions in different time points to check whether the effect may vary alone the time. The inter-layer connections are defined by the orthology information, obtained from Ensembl 84. We use the associated gene name as the unique identifier for each gene (node) in both human and mouse, and the corresponding orthologos mapping table are embedded into this two-layer network. After gene expression data pre-processing and orthologos selection, we get 19332 genes in human layer and 13656 genes in mouse layer. There are 8066 links between two layers, standing for confident orthologous mapping pairs.

As the same in single layer case, we use the binary search for parameter α in elastic net penalties to get the desired modules size for both layers. It turns out to be a grid search process in order to get two desired modules at the same time. Here we only consider the large λ_3_ in (2) which aims to find a conserved module across these two species. We use samples from all conditions to construct the basic weighted co-expression networks for two species, but different expression values under different time to define node activities. Because correlation based network construction requires as many samples while gene activities are closely related to certain conditions, including the exposed time period.

Figure 3 shows the most active module (the first identified) for human and mouse at the time point 2 hours, visualized by muxViz [8]. The conserved module is acquired when λ_3_=1000 in (2), where inter-layer links mean the orthologous gene pairs. The gene lists of two modules are attached in table file (S2.xlsx). Gene ontology enrichment analysis indicate that there are several GO terms such as response to virus (G0:0009615) and cellular response to type I interferon (G0:0071357) are significantly enriched for mouse, while GO terms like response to endoplasmic reticulum stress (G0:0034976) and topologically incorrect protein (GO:0035966) for human. Both modules shows some cellular response to topologically incorrect protein (GO:0035967) or virus (GO:0051607), which is closely related to the functional role of Th17 differentiation played in pathogenesis of autoimmune and inflammatory diseases [39].

**Figure 3:**
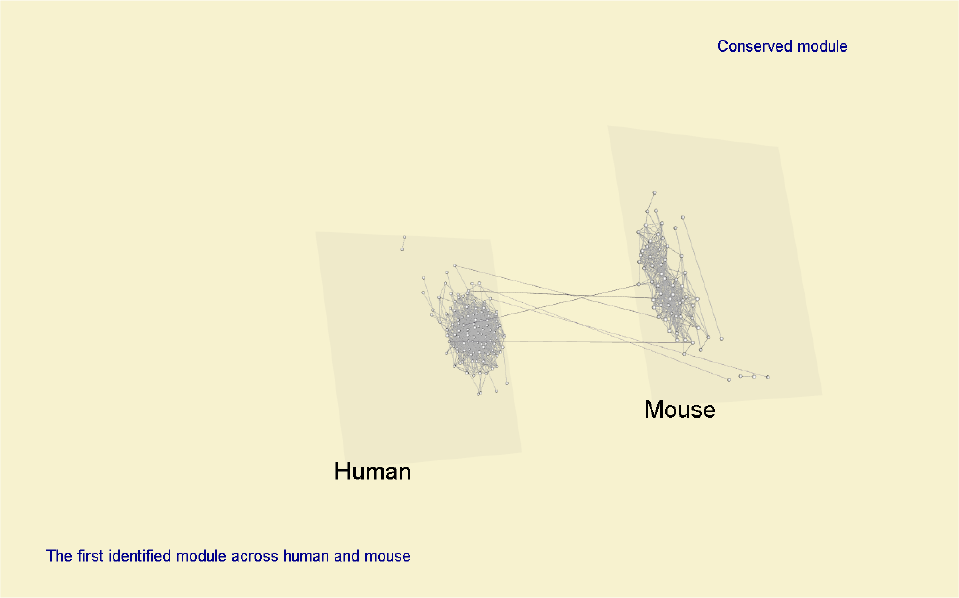
Conserved modules for human and mouse. Visualized by muxViz [8]

Besides the early stage of Th17, we also explore the characteristics of other time points (12h 24h, 48h and 72h). Figure 5 shows the most active module (the first identified) for human across these four time points. We can see that some shared genes show activity along the time and play important roles in these networks. All gene lists are attached in table file (S3.xlsx). Gene ontology enrichment analysis show that all modules are significantly related to several biological processes, which are consistent with previous studies [].

**Figure 4:**
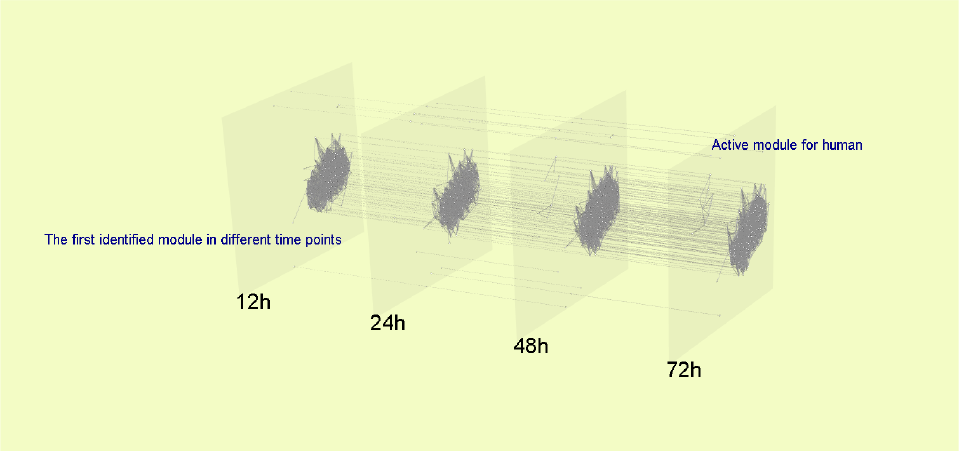
Active modules for human across time points. Visualized by muxViz [8]

The first active modules identified in each time points themselves show some differences, indicating that modification of Th17 would consistently have an impact on the cell along the time. From the algorithmic point view, when fixing λ3 in (2) we need to seek the optimal parameters in the elastic penalties for both species. i.e *f_1_*(x) = α_1_‖x‖_p_ + (1-α_1_)‖x‖2 and f (y) = α_2_‖y‖*p* + (1-α _2_)‖y‖_2_. The grid search uses a binary search for each to seek a combination of two *α*s for desired modules. Table 1 shows the results,

**Table 1:** Best parameters for equation (2)

|  | 12h | 24h | 48h | 72h |
| --- | --- | --- | --- | --- |
| $\alpha_1$ | 0.675 | 0.675 | 0.563 | 0.563 |
| $\alpha_2$ | 0.042 | 0.028 | 0.018 | 0.028 |

We use an integrative gene list enrichment analysis tool Enrichr for human [6], which does not only provide the pathway and gene ontology enrichment analysis, but also has a visualization tool with each. Figure 5 shows top GO items enriched by the most active modules different in different time points and their relationships, check supplementaty file part 3 for other time points (Figure S1-S3). Although these modules have a lot genes in common, functions enriched by the first identified module changed slightly along the time. Structural constituent of ribosome (GO:0003735) appeared frequently as a top enriched term in all time points, served as a fundamental function.

**Figure 5:**
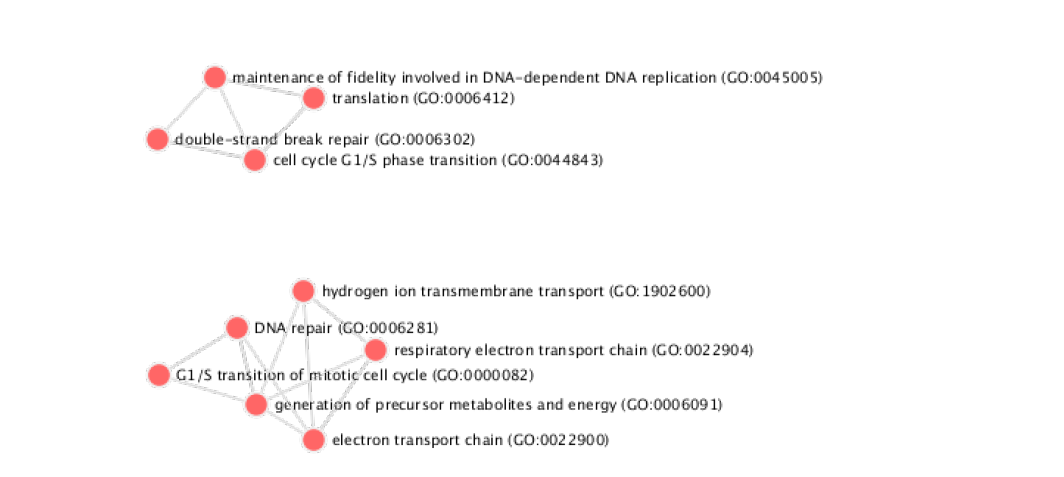
GO terms network of the identified module at 12h

### 4 Discussion and Conclusion

There have been many works discussing about key individual components such as transcription factors which play important roles in a biological process. Popular tools such as limma [35] uses statistical models to find significantly expressed genes, which almost becomes a standard pre-processing for further study. Take Th17 cell differentiation for example, previous works [36, 43, 39] reveals important transcription factors involved and related mechanism. 0ur model is designed not for individual gene detection tasks such transcription factors identification or gene prioritization, but for providing a complementary from modular perspective. Modules identified from genome-wide network may reveal system-level properties of related biological mechanism. The goal of algorithm 1 is to establish a computational approach that only uses gene expression data from different conditions, to find biological modules which show significant responses caused by expression changes.

The continuous optimization method, especially the convex optimization also offers high efficient computational tools other than widely used heuristic algorithms [19, 48] or discrete optimization [10, 13] for active modules identification. Furthermore, convex optimization methods always enjoy the guarantee with respect to running time and accuracy. On the one hand, this guarantee makes the solution more reliable even unique given precise input. On the other side, the so-called optimal solution fully relies on algorithmic input which poses a higher demand on data preprocessing and model assumption. Take the real-word data studies in section 3.2 and 3.3 for example, slight differences on how to compute node scores or edge weights may lead totally different results. Conversely the uncertainty in heuristic algorithms may offer flexibility about model assumption and algorithmic input. In other words, the gap between computational model and real case does exist. From the software design and implementation view, open source and user-friendly tools have more advantages. A large number of reliable open source libraries can be easily found for mature convex optimization techniques. And it is not difficult to implement the core part of them. In contrast, the implementation of specific purpose heuristic algorithms or integer programming is challenging.

This paper describes a general continuous optimization based active modules identification method for multilayer gene coexpression networks. With proper replacement of node (gene) similarity matrix and node activities, the proposed methods can be easily extended to other applications. The idea of formulating the conserved modules identification across multiple layers under a uniform framework can also be enriched with more sophisticated considerations, such as multiple data source fusion instead of using single gene expression profiles. Future works may be more integrated with specific applications.

## Funding

This work has been supported by the…

## Footnotes

1 http://www.yeastgenome.org/help/analyze/go-term-finder#pvalue

2 http://www.ensembl.org/

